# TSProm: Deciphering the Genomic Context of Tissue Specificity

**DOI:** 10.1101/2025.10.30.685161

**Authors:** Pallavi Surana, Pratik Dutta, Nimisha Papineni, Rekha Sathian, Zhihan Zhou, Han Liu, Ramana Davuluri

## Abstract

Characterizing tissue-specific (TSp) gene expression is crucial for understanding development and disease; however, traditional expression-based methods often overlook the latent “regulatory grammar” embedded in the non-coding DNA, particularly in distal promoter regions. Here, we introduce *<u>TSProm</u>*, a framework that specializes a DNA foundation model (DNABERT2) to decipher the long-range regulatory logic of TSp promoters at the gene isoform level. The contributions of our work are two-fold. First, we propose a novel comparative design that trains two distinct models, A: for general promoter biology and B: for TSp regulation. These models enable the precise isolation of sequence motifs around the transcription start site that uniquely define tissue identity. Second, we introduce a comprehensive explainable AI (xAI) module that integrates attention-based discovery with model-agnostic SHAP analysis to provide robust, cross-validated interpretations of learned features. Applying this framework to human brain, liver, and testis promoters, we identified and validated clinically relevant transcription factors (TFs) in the brain, including *SP1, MYC*, and *HES6*, and confirmed their known roles in diseases such as gliomas and neuroblastomas. Our analysis further revealed that *C2H2* Zinc Finger proteins are a dominant feature of the global landscape of TSp gene regulation. TSProm provides a novel and interpretable framework for identifying TSp gene regulatory elements, offering powerful computational tools for the study of tissue-specific gene regulation in normal and disease conditions.

## Introduction

Precise regulation of gene expression underlies development, homeostasis, and disease in multicellular organisms. Distinct tissues and cell types exhibit unique transcriptional programs shaped by chromatin state, transcription factor (TF) binding, and sequence context [Davuluri et al., 2008, Sonawane et al., 2017]. Disruption of these regulatory mechanisms contributes to conditions ranging from cancer to neurodegeneration and often presents in tissue/cell-type specific manner [Cavalli and Heard, 2019]. Despite extensive transcriptomic profiling, the sequence features encoding tissue-specific *(TSp)* regulation remain incompletely understood. Regulatory elements controlling TSp promoter usage play an essential role in organ development and disease dysregulation, with widespread regulation of alternative mRNA transcripts further influencing promoter activity [Davuluri et al., 2008, Carninci et al., 2006, Surana et al., 2024]. Understanding their language, taking long-range context is essential for revealing which functional modules get disrupted in complex diseases such as cancer, where promoters often lose their TSp-icity and acquire increased plasticity [Haigis et al., 2019, Frost, 2021, Sonawane et al., 2017].

Traditional approaches to characterize TSp-icity focus on differential expression or marker genes, which overlook the latent “regulatory grammar” embedded in non-coding promoter and enhancer sequences [Smith et al., 2007, Sonawane et al., 2017]. In addition, most methods focus on core and proximal promoters [regions upstream of the transcription start sites (TSS)] and not distal regions. Recent advances in transformer architectures have shown that deep learning models can extract biologically meaningful representations directly from raw DNA across both coding and non-coding genomic regions like promoters, enhancers, silencers, insulators, splice sites, and more [Consens et al., 2023, Guo et al., 2025, Zhou et al., 2024, Ji et al., 2021, Dalla-Torre et al., 2025, Fishman et al., 2025, Nguyen et al., 2023].

Utilizing the capability of these models and their significant applications in understanding alternative promoters’ regulatory regions, we introduce a gene isoform-level <u>T</u>issue <u>S</u>pecific <u>Prom</u>otor DNA Language model *<u>TSProm</u>*, extending DNABERT2, a 2nd generation transformer-based genomic sequence-only foundation model to classify and decipher TSp promoter biology. It utilizes Byte Pair Encoding (BPE) tokenization and Attention with Linear Biases (ALiBi) to efficiently process long, multi-species DNA sequences [Zhou et al., 2024, Consens et al., 2023]. We wanted to explore TSp-icity in normal tissues so that we can develop an understanding of the specific rules of normal tissues and further validate their relevance in complex genomic diseases.

We implemented TransTEx [Surana et al., 2024] to curate promoter expression groups from human available at TransTExdb and mouse. Promoters were classified into five groups: TSp, representing transcripts uniquely expressed in a single tissue; TEn, transcripts expressed in one or more but fewer than 50% of tissues; Wide, transcripts expressed in more than 50% of tissues; Low, transcripts with low expression or limited to a small subset of samples across all tissues; and Null, transcripts with no detectable expression or expressed only in a minimal number of samples across tissues. We extended regions around the TSS for the promoters grouped by tissue to capture both proximal and distal promoter interactions. For both human and mouse, we focused on three representative tissues based on the number of TSp trasncripts: testis, liver and brain for fine-tuning DNABERT2 and compared their performance with GENA-LM and Nucleotide Transformer model [Fishman et al., 2025, Zhou et al., 2024, Dalla-Torre et al., 2025].

To interpret predictions, we implemented a comprehensive attention-guided motif (short repeats with biological meaning) discovery module, cross-validated [Ji et al., 2021] with <u>SH</u>apley <u>A</u>dditive ex<u>P</u>lanations (SHAP)-based scores from game theory [Lundberg and Lee, 2017], which discovers and verifies sequence motifs most relevant to TSp-icity. This explainable AI *(xAI)* approach improves the understanding of TFs in the distal promoter and regions beyond, across tissues. The module also reveals candidate regulatory motifs consistent with known TF binding patterns. We further validate the findings with publicly available experimentally validated datasets in literature like JASPAR 2024 CORE database [Rauluseviciute et al., 2024], TF-Marker [Xu et al., 2022] and KnockTF 2.0 [Feng et al., 2024].

TSProm demonstrates that transformer-based encoder DNA Foundation Models (DFMs) can be effectively specialized to capture long-range TSp promoter logic, providing both predictive performance. The xAI module gives insights into the tokens/motifs (DNA language words) that are potentially responsible for TSp-icity of isoform-level gene regulation. This understanding helps in the identification of relevant TFs and the target genes that are dysregulated in complex multifactorial genomic diseases.

## DNA Foundation Models

DFMs are deep learning models trained with self-supervised learning on large unlabeled datasets such as genomes. By enabling attention outputs, they provide detailed maps that offer xAI into how nucleotide tokens contribute across transformer layers [Ji et al., 2021, Le et al., 2022].

We employed three state-of-the-art DFMs for sequence analysis: DNABERT-2, Nucleotide Transformer and Gena-LM. DNABERT-2, a bidirectional transformer encoder, was utilized for its superior performance in representation learning tasks. The model processes DNA sequences BPE tokenization to replace the kmer tokenization to generate contextual embeddings through masked language modeling pre-training on multi-species genomic data. The model has 12 transformer encoder layers, 12 self-attention heads, and approximately 117 million parameters. This also aligns with the semantics of DNA sequences, where information context naturally varies with sequence length. We fine-tuned DNABERT-2 using task-specific labeled datasets with a learning rate of (2e-5, 1e-5, 3e-4, 5e-6 and others) and batch size of 32 or 64 [Liu et al., 2024, Zhou et al., 2024].

Gena-LM employs an autoregressive transformer decoder architecture with 24 layers, 1024 hidden dimensions, and 16 attention heads, and was selected for superior performance in sequence understanding and attention interpretability. This model uses single nucleotide tokenization and next-token prediction objectives during pre-training [Liu et al., 2024, Fishman et al., 2025].

The Nucleotide Transformer (NT; 500M–1000G) model is a bidirectional transformer encoder based on the BERT architecture, comprising approximately 500 million parameters. It is pre-trained using a masked language modeling objective on a diverse corpus of 3,200 human genomes and 850 genomes from other species, enabling cross-species representation learning and contextual understanding of genomic sequences. Unlike previous models, NT replaces overlapping with non-overlapping k-mer tokenization [Liu et al., 2024, Dalla-Torre et al., 2025].

All three DFMs were implemented using PyTorch and fine-tuned on NVIDIA A40 GPUs with gradient accumulation steps of 1 to accommodate memory constraints.

## Datasets

We use the tissue–transcript groupings from TransTEx [Surana et al., 2024] and our custom groupings of Mouse TransTEx (Data from Li et al. [2017]). We construct a train, validation, and test split (80:10:10) of the top three TSp sites (testis, brain, and liver) together with the four remaining groups (Wide, Null, TEn, and Low) for fine-tuning. This setup was evaluated with DNABERT2 and GENA-LM. The number of unique transcripts based on the TSS along with the tissues common between human and mouse TransTEx is summarized in (**Appendix A**).

### Curated datasets with varying longer sequence lengths

We utilized transcriptomic groupings for human promoters from both the Mouse (mm39) and human (hg38) genomes. For these species, TSp promoters for the top three tissues (label 1), while promoters of the other group (label 0). We evaluated three promoter window configurations: -3 kb to +1 kb, -2 kb to +1 kb, and -1 kb to +1 kb relative to the TSS, to test core, proximal and distal promoter context on model performance. This ensures we identify additional DNA tokens/motifs that localize near TSS apart from the core and proximal promoters [FitzGerald et al., 2004, Lambert et al., 2018]. To determine biologically relevant sequence lengths, we analyzed the TFBS and histone mark profiles from ENCODE and studied the enrichment of peaks around TSS as summarized in (**Appendix B**). We found enrichment in regions 2k bp upstream and 1k bp downstream for most promoter related TFBS. This allows the models we built to capture core, proximal distal regulatory signals related to TSp-icity.

### Tackling Data redundancy

To reduce redundancy among the sequences, we applied MMseqs2 clustering using *‘mmseqs easy cluster’* with a minimum sequence identity and coverage threshold of 80% [Steinegger and Söding, 2017]. This is done to retain only representative sequences from each cluster, yielding a non-redundant set for fine-tuning.

## Methods

### Fine Tuning of DFMs

To classify TSp groups from other expression categories, we fine-tuned DFMs using LoRA to minimize memory and compute requirements from HuggingFace. GENA-LMs’s AIRI-Institute/gena-lm-bigbird-base-t2t model variant, utilizes BigBird-based sparse attention and ALiBi positional encoding to handle long DNA sequences efficiently whereas DNABERT2’s zhihan1996/DNABERT-2-117M model handles a range of short to longer lengths. These models have shown superior performance in longer sequence ranges for promoters [Liu et al., 2024, Zhou et al., 2024, Fishman et al., 2025]. To validate these benchmark results on another model, we finetune with Nucleotide Transformer’s InstaDeepAI/nucleotide-transformer-500m-1000g pre-trained on whole genome DNA sequences over 3000 diverse genomes [Dalla-Torre et al., 2025].

### Models A and B of promoter biology

To understand the language of TSp gene regulation, we created negative set against a set of human and mouse promoters with sequence lengths of upstream (1k, 2k and 3k) and downstream of 1k bp (5^*′*^→3^*′*^). We used these sequence lengths due to their distribution around the TSS sites of TF binding regions which are critical in promoter regulation **(Appendix B)**. *<u>Model B</u>* : Negative set of this binary classification was same GC content and repeats patterns from the genome [Krützfeldt et al., 2020] and *<u>Model A</u>*: where we randomly selected genomic locations outside promoter sequences, ensuring that negative and positive samples do not overlap [Fishman et al., 2025, Dalla-Torre et al., 2025] **(Figure 1: Module I)**.

**Fig. 1.**
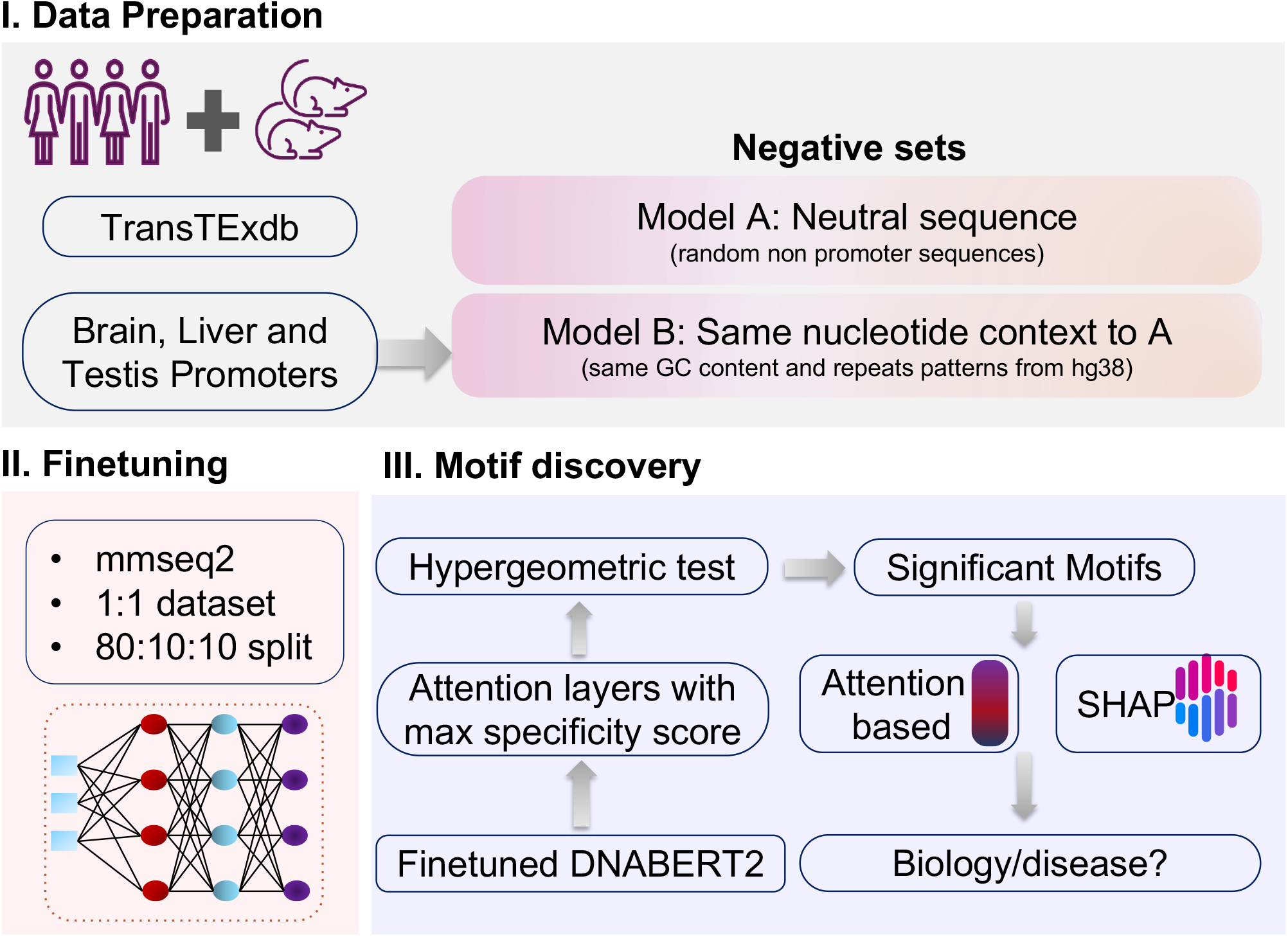
Overview of the TSProm framework. (**A**) Data preparation from multiple species and sources, including two negative sets used to construct Models A and B, respectively. (**B**) Fine-tuning using pre-trained genomic large language models (`DNABERT2, GENA-LM`). (**C**) Explainable AI and motif discovery workflow depicting attention-based interpretation and motif enrichment analysis.

### Model evaluation metrics

We evaluated our fine-tuned models using key metrics: Accuracy, F1-Score, MCC, Precision, and Recall. Accuracy measures the rate of correct predictions, F1-Score assesses precision and recall balance, MCC provides a robust correlation coefficient regardless of class size evaluates the model’s ability to distinguish between classes at various thresholds. These metrics collectively ensure a thorough assessment of precision, recall, classification quality, and discriminative capacity **(Appendix C)**.

### Attention-based motif discovery

#### Specificity scoring per layer and motif selection

We used the best fine-tuned models to extract attention-based motifs and assess their enrichment relative to the TSS **(Figure 1: Module III)**. For each sequence, token-level attention scores were obtained by averaging across all attention heads within each layer and then normalized relative to the maximum score per sequence.

Let **q**^∗^ denote the query vector corresponding to the special classification token [CLS], which is prepended to each input sequence and used for downstream prediction. Let *d* be the hidden dimension of **q**^∗^, and let **k**_*j*_ represent the key vector of the *j*-th BPE token, where *j* ∈ *{*1, …, *J}* and *J* is the number of tokens in the sequence. The attention weight assigned to token *j* across all *H* attention heads is defined as

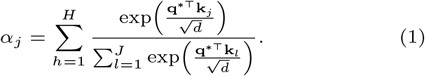

Here, *α*_*j*_ quantifies how strongly the global [CLS] representation attends to token *j*, and thus provides an attention score for sequence segments of variable length [Ji et al., 2021, Zhou et al., 2024].

Normalized token-level attention scores are defined:

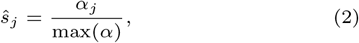

where max(*α*) is the maximum attention weight within the sequence.

### Layer specificity

To identify the most informative layers, we defined a specificity score based on the idea of z-score

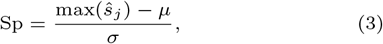

where ŝ are the normalized attention scores, and *µ, σ* denote their mean and standard deviation, respectively. Based on this metric, the middle layers (5, 6, and 7) together with the final layer were identified as the most informative across models, which were selected for motif discovery.

### Motif selection

Tokens were considered candidate motifs if they satisfied:

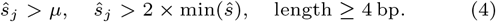

Enrichment in positive versus negative sequences was tested using the hypergeometric distribution:

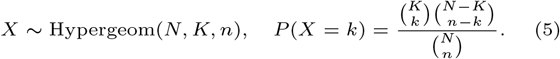

where *N* is the total number of sequences, *K* the number of positive sequences, *n* the number of motif-containing sequences, and *k* the number of positives containing the motif [Barash et al., 2001, Ji et al., 2021]. This distribution models the probability of a specific success in a sample drawn without replacement from a finite population set.

Ahocorasick algorithm was applied to [Tumeo and Villa, 2010, Ji et al., 2021] for multi-pattern matching and ensures that motifs are present at least once. False discovery rate (FDR) correction with *α* = 0.01 was applied to identify significantly enriched motifs. Motifs with fewer than three instances were discarded.

Redundant motifs were merged using gapless overlap, and *±*12 base pair (bp) sequence windows around motif centers were extracted for visualization. De novo motif discovery was performed with `MEME`, and identified motifs were compared against the JASPAR 2024 CORE database [Rauluseviciute et al., 2024] using `TOMTOM` for TF annotation [Bailey et al., 2009].

#### Validation of motifs: SHAP analysis

We applied SHAP KernelExplainer to interpret finetuned DNABERT2 predictions at the token level [Lundberg and Lee, 2017, Mosca et al., 2022]. Each promoter sequence was tokenized, and a background baseline was constructed by replacing all tokens with the model’s mask token. Using this baseline, KernelExplainer estimated SHAP values for the positive class, quantifying each token’s contribution to the classification logits [Kokalj et al., 2021]. To interpret the contribution of sequence tokens, we computed SHAP values for each token across all input sequences. For every unique token, we calculated the mean SHAP value, its absolute mean SHAP value, the frequency of occurrence, and the standard deviation. A 95% confidence interval (CI) for the mean was estimated as 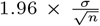, where *σ* is the standard deviation of SHAP values and *n* is the number of token occurrences. Tokens with *<* occurrences were excluded to avoid noise. Tokens were then ranked either by frequency, mean SHAP value, or absolute mean SHAP value, and the top *k* tokens were selected for visualization. Relative importance of the top positive and negative tokens mean SHAP values were plotted (Example: Figure 3D).

#### Motif positional analysis

Consensus motif positions were first identified within each input sequence. The genomic start coordinate of each motif was then computed by adjusting the sequence start or end depending on the strand orientation. Motif–TSS distances were obtained as the difference between the motif start site and TSS. To visualize positional preferences, motif–TSS distances were aggregated into 100 bp bins, and the mean attention score was calculated per bin. A three-bin rolling average (≈ 300 bp smoothing) was applied to reduce local variability. Distributions were visualized using kernel density estimation (KDE) alongside histograms, with the TSS marked at 0 on the x-axis.

### Clinical relevance of motifs

Predicted motif regions in TSp promoter sequences were validated by cross-referencing the corresponding TFs with TF-Marker [Xu et al., 2022] and KnockTF 2.0 [Feng et al., 2024] databases. TF-Marker provides experimentally verified TFs that act as cell type- or cell line-specific markers, many of which are critical for cancer cell identity and tumor heterogeneity. KnockTF 2.0 compiles transcriptomic profiles following TF perturbations, offering insights into regulatory pathways and oncogenic mechanisms controlled by them. Together, these resources enabled us to confirm both the TSp and cancer-relevant roles of the significant TFs.

## Results

### Models to understand promoter and TSp promoter biology

The TSProm model is built upon the multi-genome pre-trained DFM, DNABERT2. To adapt the pre-trained model parameters for TSp classification and interpretability of the significant tokens/motifs, we employed transfer learning techniques by using the frozen weights from the model. The dataset was prepared by implementing TransTEx which divides the transcriptome into 5 expression groups using isoform-level expression data from normal tissues [Surana et al., 2024] [Database: TransTExdb]. We use the database and implement this method on Mouse Body Map data [Li et al., 2017] to make MouseTransTEx.

We wanted to benchmark against a more recent attention-based DFM model, so we chose GENA-LM [Fishman et al., 2025] and observed similar performances on fine-tuning tasks across 2k, 3k, and 4k lengths around the TSS (Figure 2A). In fact, DNABERT2 models show slightly higher accuracy and F1 scores. Hyperparameter tuning was performed based on the learning rate, batch size, number of epochs, weight decay, and dropout rate (**Appendix D** also includes NT results not shown in the plot). We train models for each of the top three TSp sites (testis, liver, and brain) versus Null, Low, Wide, and TEn groups, and plot the best result from each tissue (**Figure 2A**). An important factor underlying the limited performance of our models may be the choice of sequence window (≈ 80% F1). Regions 1 kb downstream of the TSS include the first intron of regulatory promoters, which is underexplored due to its indirect involvement in gene regulation [Davuluri et al., 2008, Carninci et al., 2006]. Accounting for strand information is critical, and this is inherently addressed by the bidirectional nature of DFMs, as promoter complexes are bidirectional [Anderson and Anderson, 2024, Carninci et al., 2006]. We explored an alternative approach (**Appendix E**) by segregating the data into CpG and non-CpG promoter groups which do not show better performance.

**Fig. 2.**
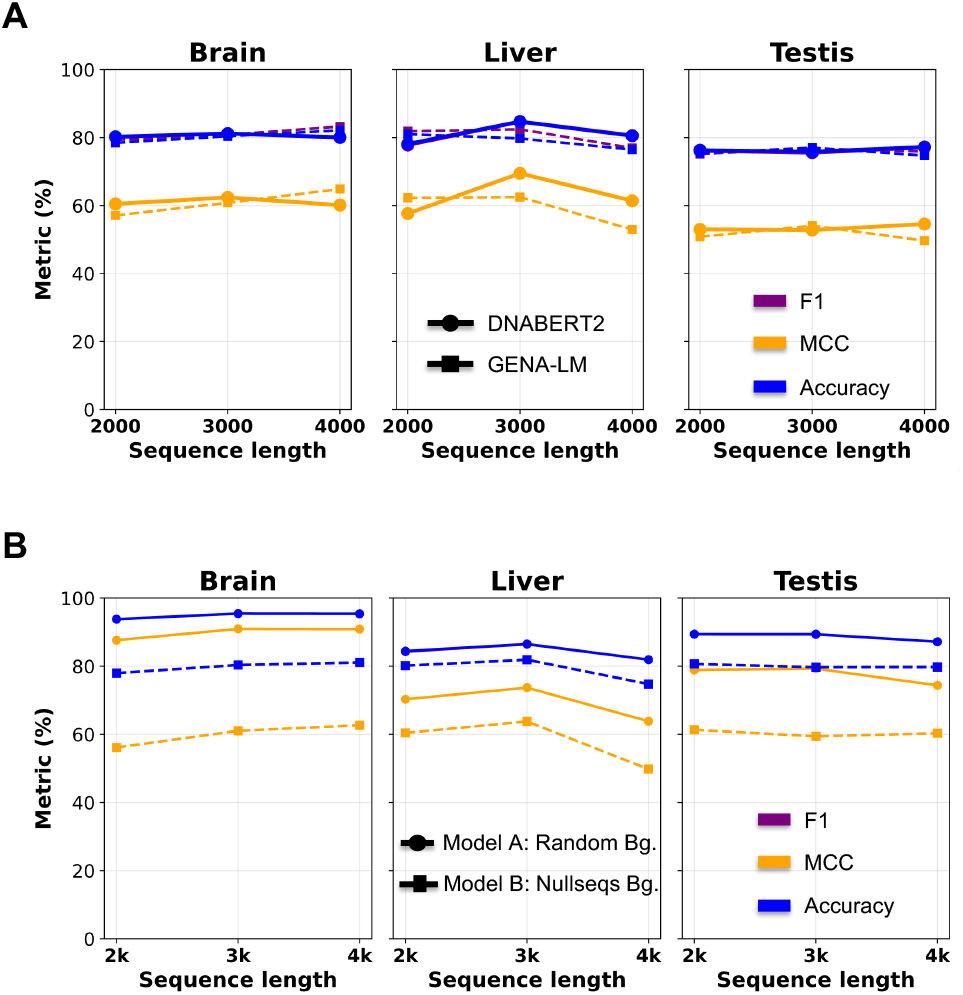
(**A**) Fine-tuning performance of `DNABERT2` and `GENA-LM` on TSp versus other expression groups across the top three tissues, showing *F*_1_, MCC, and accuracy (colored lines). (**B**) Comparative performance of Models A and B for `DNABERT2` on TSp data.

Hence, we built Model A and Model B for each tissue **(Figure 1: Module I-II)**.

1. Model A: general promoter biology
2. Model B: decipher TSp code

We observe better performance with Model A (F1 = 0.85– 0.95) as label 0 are random non-promoter sequences across the human genome (hg38), relative to Model B where label 0 are non-TSp sites matched for GC content and repeat patterns (F1 ≈ 0.8) for 3k length. Model A focuses on finding general tissue-wise promoter logic whereas Model B find TSp promoter logic highly specific to the tissue of interest. We then applied these models to the xAI motif discovery module **(Figure 1:Module III, Figure 2B)**.

### Deciphering brain regulatory language and disease relevance

We analyzed models A and B separately, and the logic here is to find the *B* ∩ *A*^*c*^ tokens/motifs, so we get the unique motifs regulating TSp-icity. We follow the attention analysis as described in *Section ‘Attention-based motif discovery’*.

In Model B, we identified 82 unique significant motifs for motif significance testing, of which 69 matched targets in the JASPAR database, including 29 from vertebrates. To ensure robustness, we extracted attention scores only from correct model predictions. Among the motifs identified, 20 corresponded to C2H2 zinc finger factors (ZNFs), 8 to basic helix–loop–helix (bHLH) factors, and 1 to a basic helix–span–helix (bHSH) factor. We also observed the “GGGGCGGGG” repeat to be significantly enriched in both Models A and B, prompting further investigation of its biological/ disease relevance. Motifs mapping to known TFs were identified, as illustrated in **Figure 3A**.

**Fig. 3.**
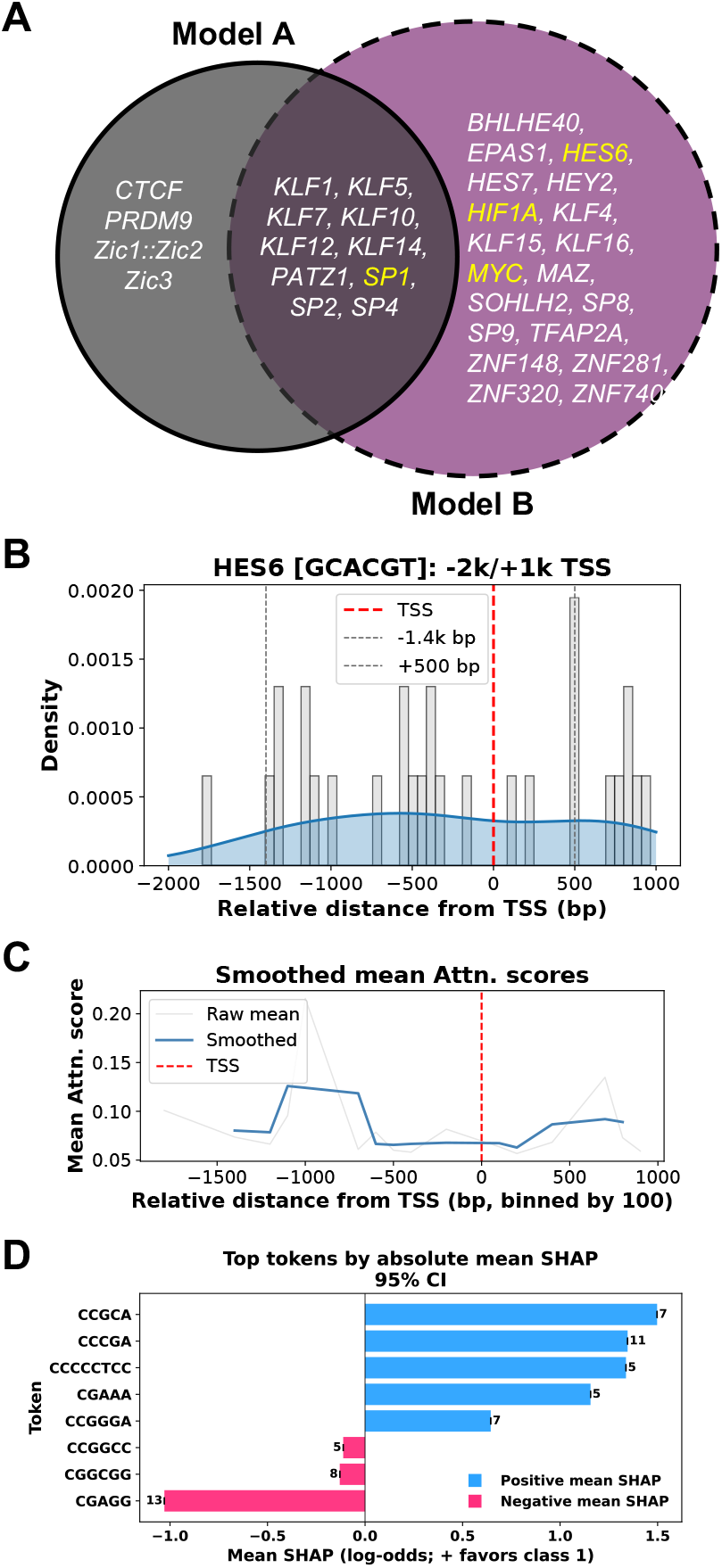
(**A**) Overlap of top significant motifs in brain identified across Models A and B found among JASPAR hits. (**B**) Density of GES6 transcription-factor motif occurrences relative to the TSS (red), with upstream −1.4 kb and downstream +500 bp regions indicated. Smoothed mean attention scores relative to the TSS, highlighting enrichment regions. (**D**) Top sequence tokens ranked by absolute mean SHAP values with 95% confidence intervals. Token counts shown above each bar.

Models A and B share 10 TF motifs from JASPAR, with *SP1* emerging as one of the experimentally validated clinical hits. It is listed in the TF Marker [Xu et al., 2022] as relevant to cancer cells in glial populations (*Appendix F3, Table 4-5*). *Appendix F2* highlights that Model B concentrates on SP1 motifs, with elevated attention and SHAP signals across the –1600 to +500 bp region, indicating its role in TSS-proximal specificity. In contrast, Model A distributes SP1 motifs more evenly, emphasizing core and proximal promoters and reflecting broader brain promoter features (**Appendix F1**).

*SP1*, a ubiquitous TF, binds to gene promoters and plays a central role in transcription. This TF has also been found to be cloned into the FLEX calling cards system for cell type–specific applications and is known to interact with gene *BRD4*. Analysis from TransTExdb shows that *SP1* expression is predominantly Wide or low transcripts [Surana et al., 2024]. It was striking that *SP1* is a significant marker of gliomas in the brain. Also, *BRD4* -mediated *MYC* TF degradation has significant translational implications [Devaiah et al., 2020]. Additionally, the *Sp1–TIMP1* axis has been reported as a potent biomarker for evaluating immune cell infiltration at tumor sites and for tracking malignant progression in glioblastoma of the brain [Liu et al., 2022]. This highlights the critical role of these TFs in normal and disease biology.

Another significant motif “GCACGT” enriched among Model *B* ∩ *A*^*c*^ motifs, maps to 2 TFs of interest, *HES6* (**Figure 3B-D**) and *MYC*, both belonging to the bHLH class of TFs. Most of *HES6* ‘s transcripts from transtex are non-TSp except *“ENST00000409356”* which is specific to the pituitary gland. **Figure 3B** shows the distribution of attention scores, which is enriched across the 3k bp window around – 1400 to 500 bp. Smoothed attention profiles show elevated attention scores in the same region, a few bp away from the TSS, suggesting the value of this model’s predictions in looking at longer regions around TSS (**Figure 3C**). SHAP analysis further focuses on similar repeats like *CGA, GC* –rich tokens, particularly *HES6* -like TFs, as key drivers of positive classification (**Appendix F2**).

According to the KnockTF 2.0 [Feng et al., 2024] database, *HES6* shows enrichment in glioblastoma cell lines, while *MYC* is enriched in neuroblastoma cell lines. (**Appendix F3**). KnockTF 2.0 provides a comprehensive dataset of TF/cofactor knockdown and knockout experiments across multiple tissues and species. Expression analysis indicates that *HES6* co– factors are significantly upregulated in low-grade gliomas and glioblastomas of TCGA dataset, underscoring their critical role in brain cancer progression. According to TransTExdb, most of the genes *HES6* targets are predominantly low or widely expressed in normal tissues. *OASL* a target gene of this TF that reprograms fatty acid metabolism to enhance oncogenesis which is also a key type-I interferon stimulated gene [Li et al., 2025].

Similarly, *MYC* TF demonstrates significant enrichment in neuroblastoma cell lines, consistent with its well-established role in neural lineage cancers. Amplification of the *MYCN* oncogene, a member of the *MYC* family, is a hallmark of high-risk pediatric neuroblastoma [Liu et al., 2021]. Moreover, *MYC* itself is activated as a potent oncogene through mechanisms such as focal amplification of distal enhancers or enhancer hijacking mediated by chromosomal translocations [Zimmerman et al., 2018].

### Validation of TSProm framework in another tissue: Testis

To validate the framework for another tissue, we focused on the testis due to the availability of data and its biological importance. We analyzed transcription factors enriched in testis-specific promoters and identified several factors that were uniquely overrepresented in this tissue, including *RFX1, RFX2, RFX3, Patz1, SP2, SP5*, and multiple members of the zinc finger (*ZNF*) family.

Among the identified factors, *RFX2* stands out as a key regulatory gene involved in spermiogenesis—the process by which spermatids mature into sperm [Kistler et al., 2015]. *RFX2* binding sites were also found to be enriched in super-enhancer regions associated with glioblastoma, suggesting that this factor plays a broader role in coordinating complex transcriptional programs beyond the testis [Sugiaman-Trapman et al., 2018]. These distal enhancer elements were effectively captured by the TSProm framework, demonstrating its ability to identify both proximal and distal sequence features that drive tissue-specific regulation (**Appendix G**).

### Language of the Global TSp-icity

Among the significant JASPAR hits for TF motifs we find most are involved in C2H2-ZNF TF domain and bHLH (**Figure 4A, Appendix H**). C2H2-ZNFs are the largest family of TFs and are relevant throughout development, gene regulation and disease biology. [Fedotova et al., 2017] quotes that, “C2H2 ZNFs, a family of proteins are the largest but poorly explored family among eukaryotic TFs”. Genes encoding this TF contain proteins that make up ∼ 40% of all human TF genes [Ding et al., 2009]. In dysregulation, these proteins regulate transcription of downstream genes involved in proliferation, apoptosis and cell invasion thereby causing cancer invasion [Jen and Wang, 2016]. According to KnockTF 2.0 we find *ZNF213, ZNF385* reported in adenocarcinoma and leukemia cell lines [Feng et al., 2024].

**Fig. 4.**
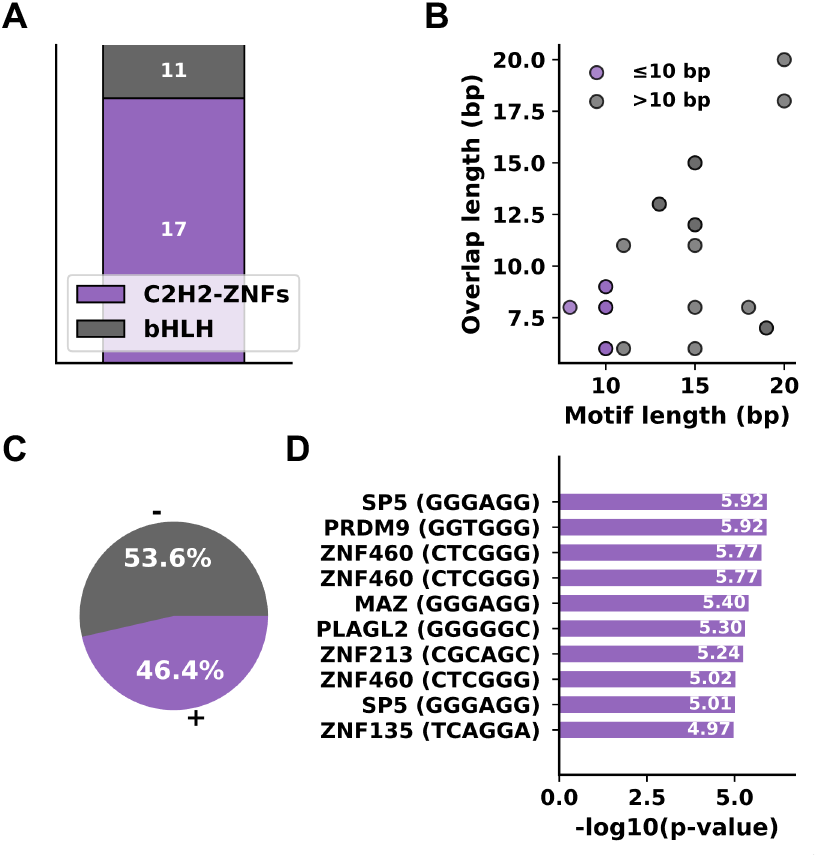
Landscape of overall TSp-icity. (**A**) Distribution of motif hits across transcription factor classes in vertebrates. (**B**) Comparison of motif overlaps grouped as short (≤ 10 bp, purple) and long (*>* 10 bp, gray). (**C**) Strandedness of motif matches. (**D**) Top enriched human motifs ranked by significance; y-axis indicates transcription factors (consensus sequences).

Similarly, we identify *MYF5* as a relevant TF Protein marker from TF-Marker, motif “ACAGCTGT” important in embryonic stem cells that can remodel chromatin in developing embryonic mesenchymal cells for myogenesis [Xu et al., 2022] (**Appendix H**).

When stratifying motifs by length, we found that longer motifs (*>* 10 bp) showed greater overlaps with target motifs than shorter motifs (≤ 10 bp) (Mann–Whitney *U* = 42.0, *p* = 0.012) (**Figure 4B**). Also, there are more negative (-) strand TFs motifs and the top human TSp TFs are summarized (**Figure 4C**). We observe *ZNF460* and *SP5* repeat in the chart, suggesting multiple significant motif patterns (**Figure 4D**).

TF *ZNF460* is significantly upregulated in gastric, colorectal cancers and promotes metastasis visa the *JAK/STAT3* pathway [Liu et al., 2023, Lan et al., 2025]. *SP5*, a C2H2 ZNF TF, modulates WNT signaling by balancing transcriptional activation and repression through distinct functional domains. Its elevated expression in hepatocellular, gastric, and colon cancers highlights its role as a context-dependent regulator of tumorigenesis [Huggins et al., 2017, Chen et al., 2006].

Overall, TSProm attention mechanism identifies significant and major TFs contributing to understanding the sequence semantics of major TFs in humans and can be extended to other species.

## Discussion

The TSProm framework represents a significant advancement in applying DFMs to understand the regulatory mechanisms modeling TSp-icity, which can then be implemented towards disease insights. By leveraging transformer architectures to analyze long-range promoter sequences up to 3kb upstream and 1kb downstream of TSS, our framework identifies clinically relevant TFs that traditional proximal and/or core promoter analyses often do not emphasize on. The framework’s ability to distinguish between general promoter biology *(Model A)* and TSp regulatory mechanisms *(Model B)* provides a systematic approach to dissect the molecular basis of healthy tissues, study its relevance in disease biology, particularly in complex diseases like cancer and neurodegeneration, where there is known TSp dysregulation of normal regulatory programs [Bianchi et al., 2020, Dos Santos et al., 2023, Sonawane et al., 2017].

Our analysis of brain-specific regulatory elements reveals key TFs with direct clinical implications for neurological diseases. The identification of *SP1* as a central regulator in brain TSp-icity is particularly significant given its established role as a biomarker for glioma and neurodegeneration progression. The *SP1-TIMP1* axis identified through our attention-based motif discovery has been experimentally validated as a predictor of immune cell infiltration and malignant progression in glioblastoma, demonstrating the translational potential of our approach. Similarly, the enrichment of *HES6* and *MYC* motifs in brain-specific promoters provides mechanistic insights into neuroblastoma and glioblastoma pathogenesis as these factors are known to be dysregulated in aggressive brain cancers. This validates TSProm’s capacity to identify regulatory elements that transition from normal tissue specification to disease-driving mechanisms. We also test the framework and validate on testis TSp regulatory landscape and identify a key regulatory of spermiogenesis which also had relevance in an aggressive brain cancer like glioblastoma [Kistler et al., 2015, Sugiaman-Trapman et al., 2018].

We find predominance of C2H2 ZNF proteins/TFs in global TSp landscape through our attention-based module. These TFs comprise ∼ 40% of the human transcriptome yet remain poorly characterized, representing possible therapeutic targets. Our identification of *ZNF460* and *SP5* as significantly enriched TF motifs, with their roles in gastric, colorectal, and hepatocellular cancers, suggests that TSp regulatory networks may be broadly dysregulated in multiple cancer types. Cross-referencing these TFs with experimentally validated databases like TF-Marker and KnockTF 2.0 provides validation for the fine-tuned models’ interpretability module. The framework’s xAI components, including attention-based motif discovery and model-agnostic SHAP analysis, ensure validations of predictions.

As DFMs continue to evolve, TSProm provides a robust pipeline for translating genomic sequences around promoters into actionable insights for precision medicine, with the potential to transform our outlook, to tackle complex genetic diseases through the lens of TSp-icity and its disruption in disease states. We further anticipate the application of this framework to other tissues, cell types contingent on data availability.

## Data and Code Availability

- TSProm: https://github.com/pallavisurana1/TSProm
- TSp Model weights: https://zenodo.org/records/17486550
- Isoform expression data: GTEx [GTEx Consortium, 2020, 2015].
- Human TransTEx groupings and code: [Surana et al., 2024], available at TransTExdb.
- Mouse BodyMap dataset: NCBI BioProject PRJNA375882 [Li et al., 2017].
- Hugging Face DNA LLMs [Fishman et al., 2025, **?**, Dalla-Torre et al., 2025]:
  - `AIRI-Institute/gena-lm-bert-large-t2t`
  - `AIRI-Institute/gena-lm-bigbird-base-t2t`
  - `jaandoui/DNABERT2-AttentionExtracted`
  - `zhihan1996/DNABERT-2-117M`

## Author contributions statement

P.S and R.D. conceived the experiment(s), P.S. conducted the experiment and analyzed the results, P.D. conducted benchmarking studies, N.P. helped with implementing fine-tuning pipeline, R.S. contributed to clinical readouts, Z.Z. and H.L. were involved in discussions attributed to their technical expertise, P.S. wrote and R.D. reviewed the manuscript.

## Acknowledgments

This work is supported in part by funds from the National Library of Medicine (NLM: # R01LM013722).

