## Supplemental Information for "TSProm: Deciphering the Genomic Context of Tissue Specificity"

#### Appendix A. Tissue choice for `TSProm`

We selected testis, brain, and liver because these tissues contained the highest number of transcripts, from which we extracted regions around the transcription start site (TSS). To address reviewer concerns about the applicability of `TSProm` to other tissues, we extended the analysis to two additional tissue transcript sets: muscle and spleen. Ovary was not included since the number of transcripts in the human data was less than 100, which was insufficient for robust training. We aimed to have at least 1,000 transcripts in the positive set to reduce the risk of overfitting for these large language models.

We used `MMseqs2` to remove sequences with more than 80% similarity, which reduced the number of unique transcripts available for each tissue. The final datasets were divided into training, validation, and test subsets to ensure balanced evaluation.

Table 1: Transcript counts by tissue and source in mouse and human `TransTEx` datasets.

| **Tissue** | **Mouse TransTEx** | **Human TransTEx** | **Human + Mouse** |
|---|---|---|---|
| Testis | 9,045 | 12,459 | 21,504 |
| Brain | 2,493 | 612 | 3,105 |
| Liver | 427 | 774 | 1,201 |
| Ovary | 1,032 | 71 | 1,103 |
| Muscle | 264 | 338 | 602 |
| Spleen | 348 | 187 | 535 |
| Lung | 242 | 142 | 384 |
| Bone marrow, blood, vessels | 156 | 204 | 360 |
| Heart | 219 | 126 | 345 |
| Adrenal gland | 150 | 117 | 267 |
| Small intestine | 143 | 57 | 200 |
| Vesicular / salivary gland | 10 | 76 | 86 |
| Stomach | 50 | 2 | 52 |
| **Total (13 common tissues)** | 14,579 | 15,165 | — |

*Totals mapped to Ensembl gene names: Mouse TransTEx: 17,320; Human TransTEx: 17,619.*

#### Appendix B. Rationale for sequence length and TSS-centered regions

Transcription factor binding sites (TFBS) located near the transcription start site (TSS) typically exert the strongest influence on transcriptional activity. These sites can act as activators or repressors depending on the transcription factor and its interaction with other cofactors. Although more distal TFBS also play roles in gene regulation, their effects are generally mediated through cooperative interactions or chromatin looping.

To determine the most biologically meaningful sequence length around the TSS, we analyzed transcription factor binding data and histone mark profiles from the ENCODE consortium. The enrichment of TFBS around the TSS indicates that most experimentally validated sites occur within approximately 3 kilobases upstream and 1 kilobase downstream of the TSS. This range represents a core promoter-proximal region where transcriptional regulation is most direct, while still encompassing some distal elements that contribute to gene-specific expression (Georgakopoulos-Soares et al., 2023; Tharakaraman et al., 2008; Veerla and Höglund, 2006).

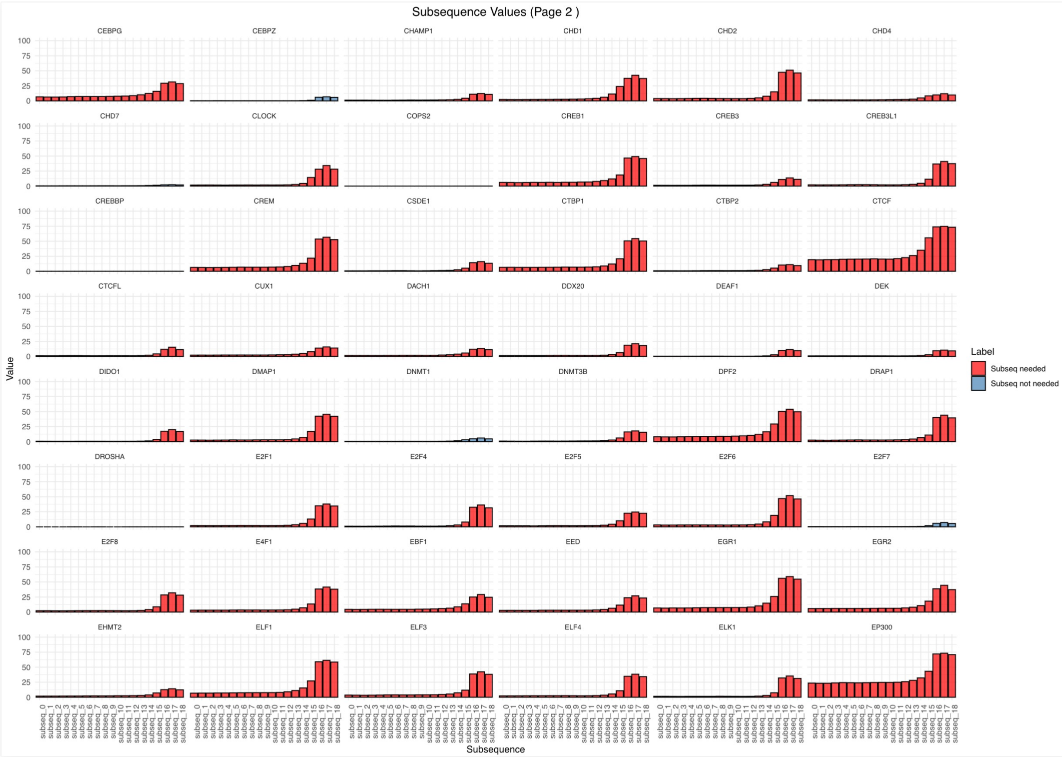


Figure 1: Illustrative example showing the enrichment of transcription factor binding sites (TFBS) near the transcription start site (TSS). The highest density occurs within ∼3 kb upstream and 1 kb downstream of the TSS, highlighting the biologically relevant region for promoter modeling.

Based on this biological evidence, we focused on modeling promoter sequences in windows extending several kilobases upstream and downstream of the TSS. This design allows us to capture both proximal promoter elements and longer-range regulatory signals.

#### Appendix C. Model Metrics

There are the summary of model metrics used to evaluate the fine-tuning of the DFMs used.

$$\text{Accuracy} = \frac{TP + TN}{TP + TN + FP + FN}$$

$$\text{Precision} = \frac{TP}{TP + FP}$$

$$\text{Recall} = \frac{TP}{TP + FN}$$

$$F1 = \frac{2 \times \text{Precision} \times \text{Recall}}{\text{Precision} + \text{Recall}}$$

$$MCC = \frac{TP \times TN - FP \times FN}{\sqrt{(TP + FP)(TP + FN)(TN + FP)(TN + FN)}}$$

#### Appendix D. Fine tuning of DNABERT2, Nucleotide Transformer and GENA-LM best models table

The table below extends Figure. 1 by reporting per–tissue results across sequence lengths (2,000, 3,000, and 4,000 bp) and learning–rate settings (see the `Run` column). The best performance was obtained when using Wide or Null as the negative class. *Wide* (widespread) promoters typically correspond to housekeeping or basal regulatory functions, whereas *Null* denotes minimal expression in adult normal tissues. These observations motivated our choice of Wide/Null as the negative set for fine-tuning, to better capture the regulatory logic of TSp promoter regions.

Unless otherwise specified, models were trained with: weight decay = $1\times10^{-2}$, warmup steps = 10, epochs = 10, and gradient accumulation steps = 1. We used sequence length in {2k, 3k, 4k} bp and learning rates in a small grid covering $[5\times10^{-6}, 7\times10^{-6}]$ and $[2\times10^{-5}, 3\times10^{-5}]$; the exact setting for each run is indicated in the table.

Table 2: Performance comparison of DNABERT2, GENA-LM, and NT (500m--1000G) models across tissues and sequence lengths. Metrics include Precision, Recall, F1-score, MCC, and Accuracy.

| Length (bp) | Tissue | TSp site | Run (LR) | Precision | Recall | $F_1$ | MCC | Accuracy | Model |
|---|---|---|---|---|---|---|---|---|---|
| 2000 | Brain | null | lr5e-6 | 80.26 | 80.23 | 80.22 | 60.49 | 80.23 | DNABERT2 |
| 3000 | Brain | null | lr3e-6 | 81.21 | 81.19 | 81.19 | 62.40 | 81.19 | DNABERT2 |
| 4000 | Brain | null | lr3e-6 | 80.07 | 80.06 | 80.06 | 60.13 | 80.06 | DNABERT2 |
| 2000 | Liver | wide | lr5e-6 | 79.57 | 78.10 | 77.82 | 57.65 | 78.10 | DNABERT2 |
| 3000 | Liver | wide | lr5e-6 | 84.77 | 84.71 | 84.70 | 69.48 | 84.71 | DNABERT2 |
| 4000 | Liver | wide | lr7e-6 | 80.83 | 80.58 | 80.54 | 61.41 | 80.58 | DNABERT2 |
| 2000 | Testis | wide | lr7e-6 | 76.73 | 76.27 | 76.16 | 52.99 | 76.27 | DNABERT2 |
| 3000 | Testis | wide | lr7e-6 | 76.97 | 75.83 | 75.56 | 52.79 | 75.83 | DNABERT2 |
| 4000 | Testis | wide | lr3e-5 | 77.37 | 77.20 | 77.16 | 54.57 | 77.20 | DNABERT2 |
| 2000 | Muscle | null | lr3e-6 | 67.97 | 67.31 | 67.00 | 35.27 | 67.31 | DNABERT2 |
| 3000 | Muscle | null | lr7e-6 | 67.97 | 67.31 | 67.00 | 35.27 | 67.31 | DNABERT2 |
| 4000 | Muscle | null | lr5e-6 | 71.15 | 71.15 | 71.15 | 42.31 | 71.15 | DNABERT2 |
| 2000 | Spleen | null | lr7e-6 | 74.48 | 73.96 | 73.82 | 48.43 | 73.96 | DNABERT2 |
| 3000 | Spleen | low | lr7e-6 | 75.05 | 72.92 | 72.33 | 47.92 | 72.92 | DNABERT2 |
| 4000 | Spleen | null | lr5e-6 | 78.31 | 77.08 | 76.83 | 55.38 | 77.08 | DNABERT2 |
| 2000 | Brain | null | lr2e-5 | 76.26 | 82.64 | 79.32 | 57.11 | 78.46 | GENA-LM |
| 3000 | Brain | null | lr2e-5 | 79.08 | 82.64 | 80.82 | 60.83 | 80.39 | GENA-LM |
| 4000 | Brain | null | lr5e-6 | 78.41 | 88.75 | 83.26 | 64.87 | 82.15 | GENA-LM |
| 2000 | Liver | wide | lr5e-6 | 78.20 | 85.95 | 81.89 | 62.29 | 80.99 | GENA-LM |
| 3000 | Liver | tenh | lr5e-6 | 72.78 | 95.04 | 82.44 | 62.50 | 79.75 | GENA-LM |
| 4000 | Liver | wide | lr5e-6 | 75.40 | 78.51 | 76.92 | 52.94 | 76.45 | GENA-LM |
| 2000 | Testis | tenh | lr2e-5 | 75.86 | 74.52 | 75.19 | 50.82 | 75.41 | GENA-LM |
| 3000 | Testis | wide | lr5e-6 | 77.70 | 75.83 | 76.75 | 54.08 | 77.03 | GENA-LM |
| 4000 | Testis | tenh | lr2e-5 | 72.47 | 79.68 | 75.91 | 49.67 | 74.71 | GENA-LM |
| 2000 | Muscle | low | lr5e-6 | 54.02 | 90.38 | 67.63 | 18.20 | 56.73 | GENA-LM |
| 3000 | Muscle | low | lr3e-4 | 50.00 | 100.00 | 66.67 | 0.00 | 50.00 | GENA-LM |
| 4000 | Muscle | low | lr2e-5 | 52.50 | 40.38 | 45.65 | 3.95 | 51.92 | GENA-LM |
| 2000 | Spleen | low | lr2e-5 | 75.61 | 64.58 | 69.66 | 44.22 | 71.88 | GENA-LM |
| 3000 | Spleen | low | lr5e-6 | 80.00 | 66.67 | 72.73 | 50.71 | 75.00 | GENA-LM |
| 4000 | Spleen | low | lr2e-5 | 57.63 | 70.83 | 63.55 | 19.26 | 59.38 | GENA-LM |
| 2000 | Brain | low | lr5e-6 | 68.14 | 66.98 | 66.44 | 35.10 | 66.98 | NT (500m–1000G) |
| 3000 | Brain | null | lr5e-6 | 71.36 | 71.21 | 71.16 | 42.57 | 71.21 | NT (500m–1000G) |
| 4000 | Brain | null | lr5e-6 | 74.39 | 74.04 | 73.95 | 48.42 | 74.04 | NT (500m–1000G) |
| 2000 | Liver | low | lr2e-5 | 62.25 | 62.22 | 62.20 | 24.47 | 62.22 | NT (500m–1000G) |
| 3000 | Liver | null | lr2e-5 | 59.63 | 59.55 | 59.47 | 19.18 | 59.55 | NT (500m–1000G) |
| 4000 | Liver | null | lr2e-5 | 62.60 | 62.50 | 62.42 | 25.10 | 62.50 | NT (500m–1000G) |
| 2000 | Muscle | low | lr3e-4 | 52.44 | 51.92 | 49.22 | 4.34 | 51.92 | NT (500m–1000G) |
| 3000 | Muscle | low | lr3e-4 | 50.00 | 50.00 | 49.33 | 0.00 | 50.00 | NT (500m–1000G) |
| 4000 | Muscle | low | lr2e-5 | 53.99 | 53.85 | 53.42 | 7.84 | 53.85 | NT (500m–1000G) |
| 2000 | Spleen | low | lr2e-5 | 60.43 | 60.42 | 60.40 | 20.85 | 60.42 | NT (500m–1000G) |
| 3000 | Spleen | low | lr2e-5 | 65.80 | 65.62 | 65.53 | 31.42 | 65.62 | NT (500m–1000G) |
| 4000 | Spleen | low | lr2e-5 | 55.21 | 55.21 | 55.20 | 10.42 | 55.21 | NT (500m–1000G) |
| 2000 | Testis | low | lr5e-6 | 53.44 | 53.25 | 52.60 | 6.69 | 53.25 | NT (500m–1000G) |
| 3000 | Testis | low | lr5e-6 | 57.04 | 56.25 | 54.99 | 13.27 | 56.25 | NT (500m–1000G) |
| 4000 | Testis | low | lr5e-6 | 59.15 | 59.00 | 58.83 | 18.15 | 59.00 | NT (500m–1000G) |

### Appendix E. Other approaches to build better models did not increase the model performance

#### E.1. Motivation

CpG and non-CpG promoter profiles differ in their sequence and regulatory properties, motivating us to test whether separating these groups could improve model performance (Landolin et al., 2010; Vavouri and Lehner, 2012). We therefore hypothesized that pretraining two `DNABERT2` models—one on CpG promoters and another on non-CpG promoters—followed by fine-tuning might better capture their distinct features. However, this split introduced strong tissue imbalances; for example, in liver the CpG subset is small ($\approx 250$ promoters) compared with non-CpG. Consequently, the effective sample sizes were much smaller than in our original setup, which likely explains the poor results we observed. It is also known that, in general, TSp regions contain more non-CpG promoters than other expression groups. These findings are consistent with prior reports linking Non-CpG promoters to TSp genes and CpG-rich promoters to housekeeping genes that are constitutively active across tissues (Saxonov et al., 2006; Vavouri and Lehner, 2012).

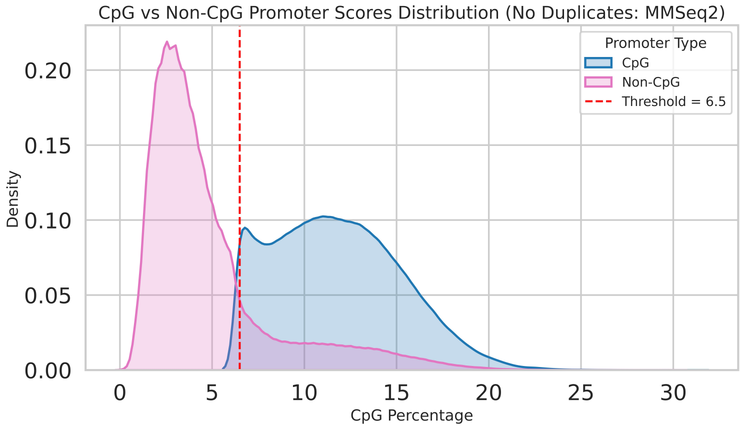


Figure 2: 201bp windows along 6k length around TSS (5k, 1k, TSS) CpG score distribution across the dataset used for pretraining.

#### E.2. Pretraining and Fine-Tuning

We pretrained models using 6 kb sequences centered around transcription start sites (TSS) from 30 species, including primates (e.g., *C. syrichta*, *A. nancymaae*, *G. gorilla*, *H. sapiens*, *M. mulatta*, *M. nemestrina*, *N. leucogenys*, *O. garnettii*, *P. abelii*, *P. anubis*, *P. paniscus*, *P. troglodytes*, *R. bieti*, *R. roxellana*, *R. norvegicus*, *S. boliviensis*) and rodents (e.g., *H. griseus*, *C. lanigera*, *I. tridecemlineatus*, *J. jaculus*, *M. murinus*, *M. caroli*, *M. musculus*, *M. pahari*, *M. spicilegus*, *M. spretus*, *N. galili*, *P. bairdii*, *R. norvegicus*, *M. fascicularis*), obtained via `biomaRt` (v2.60.1) (Smedley et al., 2009). However, during fine-tuning, the models did not show substantial improvement, as summarized in Table 3. For fine-tuning, we further stratified promoters into CpG-rich and non-CpG groups using a 6.5% CpG-content cutoff computed over sliding 201 bp windows across the TSS-centered region, but this splitting did not yield significant performance gains.

Table 3: Results after further pretraining on CpG vs. non-CpG promoters, followed by fine-tuning.

| Experiment | TSp site | TransTEx group | Length | LR | | | $F_1$ | MCC | Accuracy |
|---|---|---|---|---|---|---|---|---|---|
| *Non-CpG Further Pretrained Model* | | | | | | | | | |
| | Brain | Null | 2000 | 5e-6 | 82.95 | 82.89 | 82.83 | 65.84 | 82.83 |
| | Liver | Wide | 3000 | 3e-6 | 82.12 | 82.05 | 81.94 | 64.17 | 81.94 |
| | Liver | Wide | 3000 | 5e-6 | 81.61 | 81.57 | 81.48 | 63.18 | 81.48 |
| | Liver | Wide | 3000 | 7e-6 | 81.47 | 81.49 | 81.48 | 62.96 | 81.48 |
| | Brain | Null | 2000 | 7e-6 | 81.14 | 81.08 | 81.02 | 62.22 | 81.02 |
| *CpG Further Pretrained Model* | | | | | | | | | |
| | Brain | Null | 4000 | 5e-6 | 79.10 | 78.60 | 78.77 | 57.70 | 79.15 |
| | Brain | Null | 4000 | 7e-6 | 77.65 | 77.17 | 77.33 | 54.82 | 77.74 |
| | Brain | Wide | 4000 | 7e-6 | 76.56 | 76.50 | 76.52 | 53.05 | 76.68 |

### Appendix F. Brain Models to understand promoter biology

#### F.1. Model A for Brain

This shows that Model A shows a more evenly distribution of the identified motifs around TSS with peaks around the core and proximal promoter regions for TF Zic3 that is uniquely present in this model and not Model B. This shows our model is able to capture general promoter as well as TSp promoter enriched tokens which imply critical role in biology.

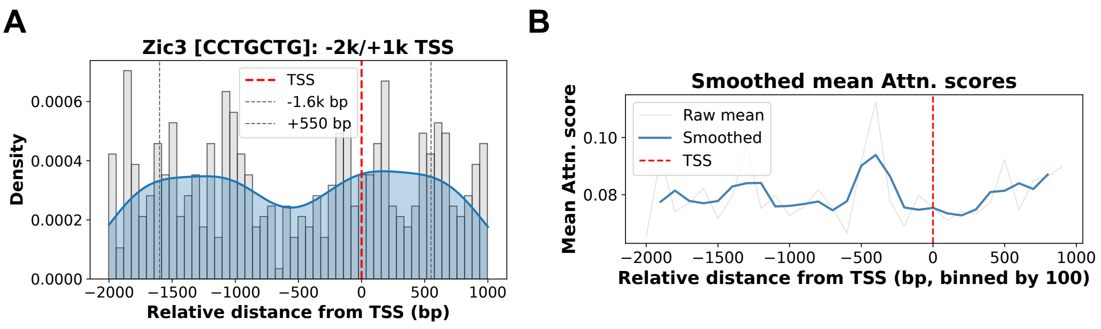


Figure 3: Detailed analysis of the Zic3 transcription factor motif. (**A**) Density of motif occurrences relative to the transcription start site (TSS), with the TSS (red dashed line), upstream −1.6 kb, and downstream +550 bp indicated. (**B**) Smoothed mean attention scores relative to the TSS, highlighting enrichment near promoter regions.

#### F.2. Model B for Brain

This shows that Model B shows a more evenly distribution of the identified motifs around TSS with peaks around the core and proximal promoter regions for TF SP1 that is ubiquitously present TF in both Models A and B. This shows our model is able to capture global brain promoter logic.

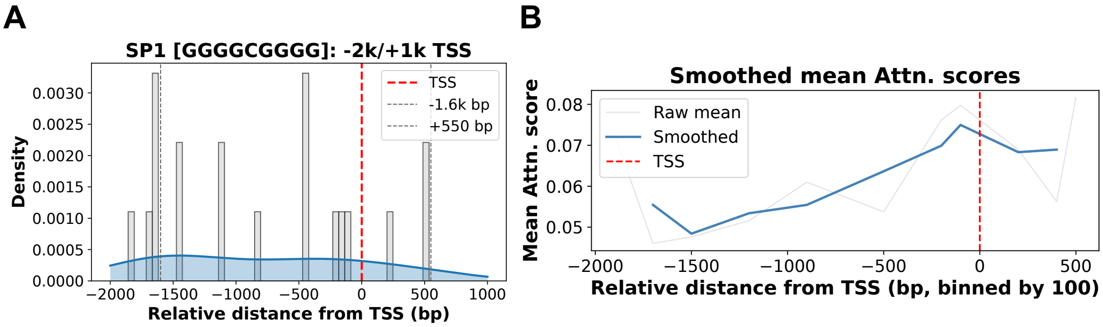


Figure 4: Detailed analysis of the SP1 transcription factor motif. (**A**) Density of SP1 TF motif occurrences relative to TSS (red), upstream −1.6 kb, and downstream +550 bp indicated. (**B**) Smoothed mean attention scores relative to the TSS, highlighting enrichment regions.

##### F.3. Model A and Model B clinically relevant TFs

We summarize the clinically relevant motifs for brain from KNOCK-TF 2.0 and TF-marker for brain specific TFs.

Table 4: Representative hits from KnockTF 2.0 (Feng et al., 2024) showing enrichment of "GCACGT" motif in glioblastoma and neuroblastoma cell lines.

| Query_ID | Target_ID | Offset | p-value | E-value | q-value | Overlap | TF_name | Tissue Type | Cell line |
|---|---|---|---|---|---|---|---|---|---|
| GCACGT | MA1493.1 | 1 | 7.46E-05 | 0.175107 | 0.0173884 | 6 | HES6 | Brain | A172 (glioblastoma) |
| GCACGT | MA1493.1 | 1 | 7.46E-05 | 0.175107 | 0.0173884 | 6 | HES6 | Brain | LN405 (glioblastoma) |
| GCACGT | MA0147.4 | 0 | 2.37E-04 | 0.556926 | 0.0291687 | 6 | MYC | Brain | BE(2)-C (neuroblastoma) |
| GCACGT | MA0147.4 | 0 | 2.37E-04 | 0.556926 | 0.0291687 | 6 | MYC | Brain | NB1691 (neuroblastoma) |
| GCACGT | MA0147.4 | 0 | 2.37E-04 | 0.556926 | 0.0291687 | 6 | MYC | Brain | BE(2)-C (neuroblastoma) |
| GCACGT | MA0147.4 | 0 | 2.37E-04 | 0.556926 | 0.0291687 | 6 | MYC | Brain | BE(2)-C (neuroblastoma) |

Table 5: Representative TF-Marker (Xu et al., 2022) hits showing motif mapping to *SP1* and *HIF1A* in brain cell types.

| Query_ID | Target_ID | Offset | p-value | q-value | Overlap | TF_name | Tissue Type | Cell type | Gene Name |
|---|---|---|---|---|---|---|---|---|---|
| RGYCWVGGGGVGGGG | MA0079.5 | -6 | 3.18E-06 | 0.00293195 | 9 | SP1 | Brain | Nerve cell (normal) | SP1 |
| GCACGT | MA1106.2 | 0 | 2.63E-05 | 0.0173884 | 6 | HIF1A | Brain | Glial cell (cancer) | HIF1A |

### Appendix G. Testis-specific regulatory language

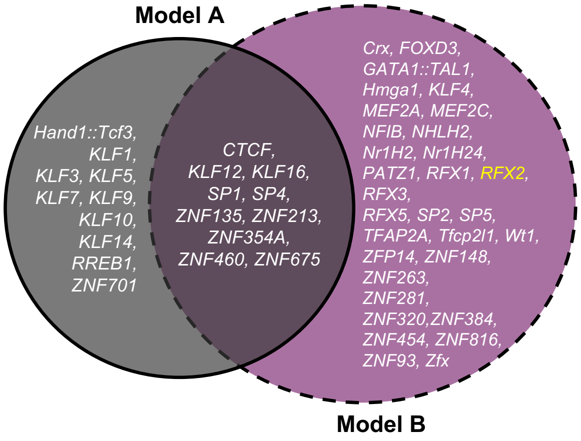


Figure 5: Overlap of top significant motifs in brain identified across Models A and B found among JASPAR hits. The same framework was applied to testis data to reveal transcription factors uniquely enriched in testis-specific promoters.

### Appendix H. High-Attention Regions Reveal Significant Global TSp Regulatory Motifs

#### H.1. Model B for all TSp promoters in human

We summarize the significant top motifs identified from the JASPAR database among the significant motifs identified in all TSp promoters in human. We identify rich "CG" repeat.

Table 6: Representative significant motifs identified from enrichment analysis from the JASPAR databse (Rauluseviciute et al., 2024).

| Query_ID | Target_ID | Offset | p-value | q-value | Overlap | Query_consensus | Target_consensus | Orientation | TF_name | Class | Family |
|---|---|---|---|---|---|---|---|---|---|---|---|
| TGCAGCTGTG | MA0521.3 | -2 | 4.50E-07 | 0.00052 | 6 | TGCAGCTGTG | CAGCTG | + | Tcf12 | bHLH | E2A |
| TGCAGCTGTG | MA0633.3 | -2 | 6.24E-06 | 0.00206 | 6 | TGCAGCTGTG | CAGCTG | + | Twist2 | bHLH | Tal-related |
| TGCAGCTGTG | MA1993.2 | -2 | 6.24E-06 | 0.00206 | 6 | TGCAGCTGTG | CAGCTG | + | Neurod2 | bHLH | Tal-related |
| TGCAGCTGTG | MA1997.2 | -2 | 6.24E-06 | 0.00206 | 6 | TGCAGCTGTG | CAGCTG | + | Olig2 | bHLH | Tal-related |
| GGAGGCYGAGGYDGS | MA1596.1 | 4 | 1.71E-06 | 0.00456 | 12 | GGAGGCCGAGGTGGG | CTCGGGAGGCTGAGGC | - | ZNF460 | C2H2 | Zinc fingers |
| GGGAGGCWGAA | MA1965.2 | 0 | 1.19E-06 | 0.00559 | 6 | GGGAGGCAGAA | GGGAGG | - | SP5 | C2H2 | Kruppel-related |
| GRTGRGGRAAAGGRGGVWGG | MA1723.2 | -2 | 1.21E-06 | 0.00566 | 18 | GATGAGGGAAAGGAGGAAGG | GGTGGGCAGGGAGGAAGCAG | + | PRDM9 | C2H2 | Multiple ZFs |
| TGCAGCTGTG | MA1635.2 | -2 | 2.17E-05 | 0.00626 | 6 | TGCAGCTGTG | CAGCTG | + | BHLHE22 | bHLH | Tal-related |
| CCGGGAGGYGGAG | MA1596.1 | 1 | 1.71E-06 | 0.00801 | 13 | CCGGGAGGCGGAG | CTCGGGAGGCTGAGGC | - | ZNF460 | C2H2 | Zinc fingers |
| GGGGGCCC | MA1548.2 | 0 | 4.98E-06 | 0.01154 | 8 | GGGGGCCC | GGGGGCCC | - | PLAGL2 | C2H2 | Zinc fingers |
| TGCAGCTGTG | MA1641.2 | -1 | 6.22E-05 | 0.01487 | 8 | TGCAGCTGTG | ACAGCTGT | + | MYF5 | bHLH | MyoD/ASC |
| TGCAGCTGTG | MA1472.3 | -1 | 6.76E-05 | 0.01487 | 8 | TGCAGCTGTG | ACAGCTGT | + | Bhlha15 | bHLH | Tal-related |
| TGCAGCTGTG | MA0048.3 | -1 | 6.39E-05 | 0.01487 | 9 | TGCAGCTGTG | GCAGCTGCG | - | NHLH1 | bHLH | Tal-related |
| TGCAGCTGTG | MA0500.3 | -1 | 8.00E-05 | 0.01546 | 8 | TGCAGCTGTG | GCAGCTGC | + | MYOG | bHLH | MyoD/ASC |
| DYYYDGYTAAWWTWTKTTTT | MA1978.2 | 0 | 1.42E-05 | 0.01659 | 20 | TTCTAGCTAATTTTTTTTTT | AATTAGTCCATTTATATTTA | - | ZNF354A | C2H2 | Zinc fingers |
| WRGGAGGSGMYGGGCYGG | MA1522.2 | -1 | 3.94E-06 | 0.01846 | 8 | AAGGAGGGGATGGGCTGG | GGGAGGGG | - | MAZ | C2H2 | Zinc fingers |
| TGCAGCTGTG | MA1619.2 | -1 | 1.25E-04 | 0.02219 | 8 | TGCAGCTGTG | ACAGCTGT | + | Ptf1A | bHLH | Tal-related |
| GGGAGGCWGAA | MA1596.1 | 3 | 9.49E-06 | 0.02228 | 11 | GGGAGGCAGAA | CTCGGGAGGCTGAGGC | - | ZNF460 | C2H2 | Zinc fingers |
| CCGGGAGGYGGAG | MA1587.1 | 0 | 1.06E-05 | 0.02490 | 13 | CCGGGAGGCGGAG | TCAGGAGGTCGAGG | - | ZNF135 | C2H2 | Zinc fingers |
| WCCAGCCTGGGCRAC | MA2121.1 | 0 | 5.69E-06 | 0.02671 | 12 | TCCAGCCTGGGCAAC | CGCAGCCTGGGC | - | ZNF213 | C2H2 | Zinc fingers |
| AGCTAGACCTCCCTGCTGC | MA1628.2 | -11 | 6.13E-06 | 0.02874 | 7 | AGCTAGACCTCCCTGCTGC | CCTGCTG | - | Zic1::Zic2 | C2H2 | Zinc fingers |
| AGCTAGACCTCCCTGCTGC | MA0697.3 | -11 | 1.23E-05 | 0.02884 | 7 | AGCTAGACCTCCCTGCTGC | CCTGCTG | - | Zic3 | C2H2 | Zinc fingers |
| GGAGGCYGAGGYDGS | MA1972.1 | 0 | 8.56E-05 | 0.03985 | 15 | GGAGGCCGAGGTGGG | GGAGGCACTGGAATG | + | ZFP14 | C2H2 | Zinc fingers |
| GGAGGCYGAGGYDGS | MA1723.2 | 0 | 7.91E-05 | 0.03985 | 15 | GGAGGCCGAGGTGGG | GGTGGGCAGGGAGGAAGCAG | + | PRDM9 | C2H2 | Zinc fingers |
| TTTTTKTWTTTTTWK | MA1125.2 | -5 | 6.49E-05 | 0.04132 | 8 | TTTTTTTTTTTTTTG | TTTTTTTT | - | ZNF384 | C2H2 | Zinc fingers |
| GGAGGCYGAGGYDGS | MA1587.1 | 3 | 9.98E-05 | 0.04223 | 11 | GGAGGCCGAGGTGGG | TCAGGAGGTCGAGG | - | ZNF135 | C2H2 | Zinc fingers |
| TGCAGCTGTG | MA0499.3 | -1 | 2.89E-04 | 0.04456 | 9 | TGCAGCTGTG | GCACCTGTC | + | MYOD1 | bHLH | MyoD/ASC |
| CAGCWMYTYGGGAGG | MA1965.2 | -9 | 9.80E-06 | 0.04599 | 6 | CAGCTACTTGGGAGG | GGGAGG | - | SP5 | C2H2 | Kruppel-related |
